# Early-life environmental effects on mitochondrial aerobic metabolism: an experimental brood size manipulation in wild great tits

**DOI:** 10.1101/2023.04.06.535828

**Authors:** Nina Cossin-Sevrin, Antoine Stier, Mikaela Hukkanen, Sandrine Zahn, Vincent A. Viblanc, Katja Anttila, Suvi Ruuskanen

## Abstract

Parental care (including postnatal provisioning) is a major component of the offspring’s early-life environment. In avian species, the number of chicks in the nest and subsequent sibling competition for food are known to affect chick’s growth, leading in some cases to long-lasting effects for the offspring. Because of its central role in converting energy, variation in the offspring’s mitochondrial metabolism could be an important pathway underlying variation in growth patterns. Here, we performed a brood size manipulation in great tits (*Parus major*) to unravel its impact on offspring’s mitochondrial metabolism and reactive oxygen species (ROS) production in red blood cells. We investigated the effects of brood size on chicks’ growth and survival, and tested for long-lasting effects on juvenile mitochondrial metabolism and phenotype. As expected, chicks raised in reduced broods had a higher body mass compared to enlarged and control groups. However, mitochondrial metabolism and ROS production were not significantly affected by the treatment either at chick or juvenile stages. Chicks in very small broods were smaller in size and had higher mitochondrial metabolic rates. The nest of rearing has a significant effect on nestling mitochondrial metabolism, yet variation in mitochondrial metabolism at the early-life stages are not associated with survival chances. The contribution of the rearing environment in determining offspring mitochondrial metabolism emphasizes the plasticity of mitochondrial metabolism in changing environments. Further studies would be needed to closely investigate what are the major environmental cues affecting the offspring mitochondrial metabolism during the growth period.

## Introduction

Parents may have the capacity to shape offspring phenotypes by influencing the offspring’s environment during development. This phenomenon, referred to as parental effects, is an important influence on offspring phenotype (Badyaev & Uller, 2009; Mousseau & Fox, 1998; Wolf & Wade, 2009). From an evolutionary perspective, parental effects, in general, are thought to improve offspring survival, growth and / or quality, hence improving parental fitness (Bonduriansky & Crean, 2018; Mousseau & Fox, 1998; Yin et al., 2019). However, it is unclear whether parental effects are always adaptive (Bonduriansky & Crean, 2018; Burgess & Marshall, 2014; Marshall & Uller, 2007; Sánchez-Tójar et al., 2020; Uller, 2008; Uller et al., 2013; Yin et al., 2019).

Parental care (e.g., postnatal provisioning) is an important early-life influence affecting offspring phenotype (Uller, 2008). For dependent offspring relying on parents to survive, it is now well established that a deficit in parental care can lead to detrimental long-term consequences (e.g., Developmental Origins of Health and Disease hypothesis), but the mechanism underlying long-lasting effects of early-life environmental conditions on offspring phenotype are not well understood (Gluckman et al., 2007; Hoogland & Ploeger, 2022; Meunier et al., 2022; Rogers & Bales, 2019).

In avian species, variation in early-life nutritional conditions and sibling competition have been widely tested by manipulating brood size (enlarging or reducing brood size) with the aim to simulate increased or reduced parental effort, thereby modulating postnatal parental care and assessing the consequences on offspring phenotype and survival. In great tits (*Parus major*), offspring from enlarged broods exhibit decreased body mass and size (wing or tarsus length) at fledging, and decreased recapture probability over the long-term, i.e. a few months after fledging (in zebra finches: De Kogel, 1997; in great tits: Hõrak, 2003; Rytkönen & Orell, 2001; Smith et al., 1989). Studies on zebra finches (*Taeniopygia guttata*) reported long-lasting effects of early-life nutritional deficits on fitness related traits, including laying initiation and breaks, hatching success, plasma antioxidant levels and flight performances (Blount et al., 2003, 2006; Criscuolo et al., 2011). Yet, the mechanisms driving the effects of early-life environmental variation (including postnatal provisioning) on the offspring phenotype and survival remain poorly understood.

Variation in metabolic rate represents one important candidate pathway underlying variation in growth patterns as it could be involved in energy allocation processes and is thought to be associated with individual fitness (Brown et al., 2018; Burger et al., 2019, 2021). Beside nestling body mass and size, several studies examined the impacts of brood size on offspring metabolic rate. In tree swallows (*Tachycineta bicolor*), nestlings from enlarged broods had 15% lower resting metabolic rate compared to individuals from reduced broods (Burness et al., 2000). On the contrary, zebra finches raised in large broods had a 9% higher standard metabolic rate at 1-year old compared to birds reared in small broods (Verhulst et al., 2006). While the association between whole-organism metabolic rate has been extensively studied to test the association between a physiological trait and fitness (or proximate traits when fitness cannot be assessed directly, see precautions here: Arnold et al., 2021; Pettersen et al., 2018), only more recently studies have focused on mitochondrial aerobic metabolism (Ballard & Pichaud, 2014; Heine & Hood, 2020; Koch et al., 2021). Studying mitochondrial respiration could reveal the cellular metabolic consequences of brood size manipulation (and thus, how variation of nutritional conditions and sibling competition influence offspring). Increased competition might have significant effect on mitochondria since organisms relying on aerobic metabolism use nutrients and oxygen for producing ATP via a set of metabolic reactions, part of them occurring within mitochondria. ATP production in mitochondria is also associated with constitutive release of damaging sub-products (e.g., reactive oxygen species, ROS), which may lead to oxidative damage that impair protein and lipid structures and promote DNA mutations (Lane, 2011; Mazat et al., 2020; Monaghan et al., 2009; Sastre et al., 2003). Thus, measuring both oxidative phosphorylation (leading to ATP synthesis) and mitochondrial ROS production (byproducts of cellular respiration) allows us to evaluate metabolic constraints and trade-offs at the cellular level (Koch et al., 2021). The efficiency by which mitochondria are able to convert ATP from a fixed amount of substrates and the determinants of this efficiency are challenging to understand as the efficiency varies between species, but also within individuals of the same species, according to age, condition and tissue (Cossin-Sevrin et al., 2022; Koch et al., 2021; Salmón et al., 2022; Stier et al., 2019, 2022).

Recent studies have found that early-life environmental stressors might impair mitochondrial function (Gyllenhammer et al., 2020; Zitkovsky et al., 2021). For example food restriction was shown to decrease basal metabolic rate in adult chinese bulbul (*Pycnonotus sinensis*) and silky starlings (*Sturnus sericeus*), and to decrease levels of mitochondrial state 4 respiration in the liver for both species (Mao et al., 2019; Zhang et al., 2018). Yet, the impact of early-life conditions on mitochondrial function and the long-lasting effects remain poorly understood.

Here, we experimentally manipulated brood size in wild great tits to test how rearing conditions (altered sibling competition for food and potential change in food availability/quality) affect nestling red blood cell mitochondrial metabolic phenotype: a promising proxy of individual performance. We aimed to test i) if brood size was important in determining nestling mitochondrial metabolism traits and associated ROS production, ii) differences in nestling growth trajectories, and if these were associated with differences in mitochondrial metabolic rates; iii) if differences in mitochondrial metabolic rates affected offspring future survival. We further iv) tested if early-life determination of mitochondrial aerobic metabolism could affect adult phenotype with potential medium-term costs (e.g., consequences on juvenile mitochondrial metabolic rates and ROS production). Finally, our experimental design allowed assessing v) the relative contributions of the foster rearing environment (from 2 to 14 days post-hatching) *vs.* the combination of genetic background, prenatal effects and early-stage rearing conditions (until 2 days post-hatching) on offspring mitochondrial metabolism. To test the impact of brood size manipulation treatment on postnatal parental care, we recorded parental feeding rates on a subsample of nests. We predicted nestlings raised in enlarged broods to have a lower body mass and size compared to control and reduced brood size. According to prior literature, the offspring mitochondrial function is sensitive to postnatal environmental conditions. In rodent models, chronic stress exposure and separation from mother during lactation led in most of the cases to a decrease in mitochondrial complexes activities and increase of ROS production (Picard & McEwen, 2018; Zitkovsky et al., 2021). We may therefore expect an enlargement of the brood size and its associated consequences, such as a decreased in parental feeding rates, to create a stressful environment leading to a general decrease of the offspring mitochondrial metabolism and increase of ROS production. Nevertheless, most of the work assessing how stressful early-life environment may impair mitochondrial function have been so far realized on mammals and the consequences in avian species and long-term effects remain elusive. Here we test the importance of brood size as a proxy to early-life environmental rearing conditions in shaping nestling mitochondrial metabolic rates, associated ROS production and later growth and survival patterns.

## Material and Methods

### a) Field site and population monitoring

This study was conducted on Ruissalo Island, Finland (60°26.055′ N, 22°10.391′ E), in a Great tit population (*Parus major* Linnaeus 1758) breeding in artificial nest boxes (n = 588 nest boxes). In Great tit, the average clutch size varies from 7 to 12 eggs (Perrins and McCleery, 1989) and the nestling period lasts from 16 to 22 days. Data for our experiment were collected during the 2020 breeding season (April to July) and during the autumn of 2020 (October to November). We monitored the breeding season progress by checking the occupation of nest boxes by great tits once a week. Clutch size, hatching date (± 24h) and fledging success were recorded.

### b) Experimental manipulation of brood size

To investigate the effects of the brood size on nestling mitochondrial function, growth pattern and subsequent survival, we performed a brood size manipulation experiment, including cross-fostering (Fig.1). We selected two nests (nest-pairs) having the same hatching date (± 24h) and conducted the brood size manipulation and cross-fostering 2 days after hatching. The initial brood size (i.e., before the manipulation) of each nest was recorded, with an average (± SEM) of 7.98 ± 0.07 nestlings per nest (ranging from 4 to 11 nestlings, n = 70 nests). Approximately half of the brood was cross-fostered between nest-pairs in order to assess the influence of the nest of origin (representing the contribution of genetic background, prenatal and early postnatal parental effects) *vs.* the nest of experimental cross-fostering (i.e., nest of rearing). The nest of rearing here reflects postnatal environmental conditions and parental effects from 2 days after hatching until fledging. The experimental design consisted of 3 treatment groups: i) a control group (C) where half of the brood was cross-fostered between nest-pairs without modifying brood size (n = 20 nests), ii) a reduced group (R) where half of the brood was cross-fostered between nest-pairs and 2 nestlings were removed from the brood (n = 25 nests), and iii) an enlarged group (E) where half of the brood was cross-fostered between nest-pairs and 2 nestlings were added to the brood (n = 25 nests) (Fig.1).

**Fig. 1.**
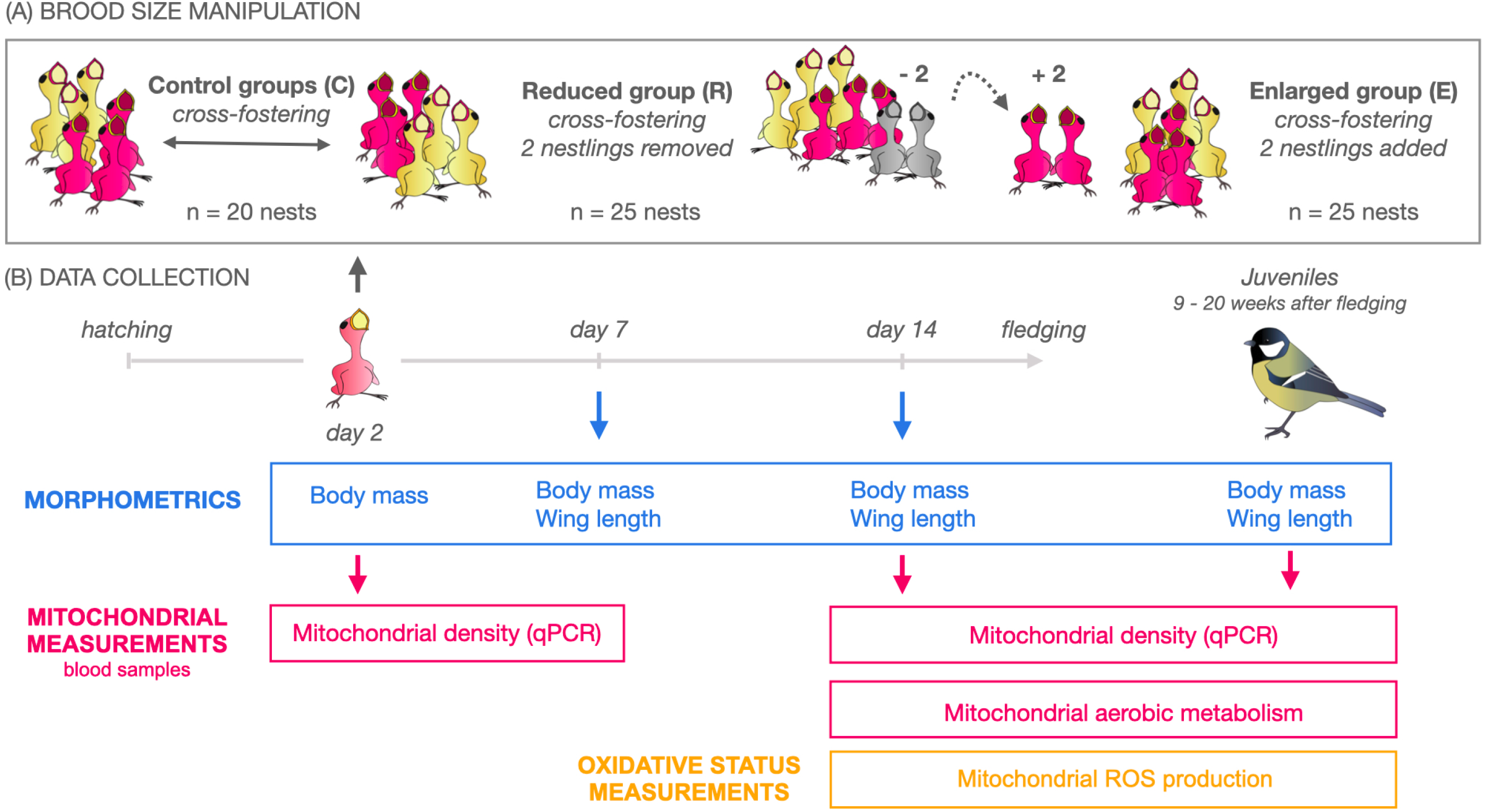
Experimental design of the study presenting the brood size manipulation (A) and collection of the data (B). Sample sizes are presented according to treatment groups: control (C), reduced (R), and enlarged broods (E). The timing of different measurements and analyses are indicated below the time-line (see Methods for details).

In total, this study included 70 great tit nests resulting in 540 nestlings monitored (*n_C_* = 150, *n_E_* = 236, *n_R_* = 154), of which 227 individuals were cross-fostered and 399 fledged (*n_C_* = 98, *n_E_* = 188, *n_R_* = 113) (see sample sizes for different measurements in Table 1).

**Table 1.** Sample-sizes according to nestling age, treatment group (C: control broods, E: enlarged broods, R: reduced broods) and the different traits measured throughout this study. The number of nests is indicated in brackets.

| Measurements | Day 2 | Day 7 | Day 14 | Juveniles |
| --- | --- | --- | --- | --- |
| Body mass/size | $n_R = 154$ (25)<br>$n_C = 150$ (20)<br>$n_E = 236$ (25) | $n_R = 121$ (21)<br>$n_C = 105$ (16)<br>$n_E = 194$ (21) | $n_R = 115$ (21)<br>$n_C = 99$ (16)<br>$n_E = 189$ (21) | $n_R = 14$ (10)<br>$n_C = 22$ (9)<br>$n_E = 31$ (15) |
| Mitochondrial DNA copy number (i.e. proxy of mitochondrial density) | $n_R = 17$ (6)<br>$n_C = 38$ (10)<br>$n_E = 16$ (5) | | $n_R = 48$ (20)<br>$n_C = 46$ (16)<br>$n_E = 55$ (21) | $n_R = 12$ (8)<br>$n_C = 16$ (9)<br>$n_E = 28$ (15) |
| Mitochondrial aerobic metabolism | | | $n_R = 35$ (19)<br>$n_C = 26$ (14)<br>$n_E = 41$ (21) | $n_R = 12$ (8)<br>$n_C = 16$ (9)<br>$n_E = 26$ (15) |
| ROS production measurements | | | $n_R = 34$ (18)<br>$n_C = 23$ (14)<br>$n_E = 37$ (20) | $n_R = 11$ (8)<br>$n_C = 16$ (9)<br>$n_E = 26$ (15) |

Before the brood size manipulation, nestlings from nest-pairs were weighed on an electronic scale (body mass ± 0.1g) and individually marked (nail-clipping). We performed blood sampling on a subsample of nestlings 2 days after hatching (1 - 10µL from the tarsus vein using heparinized capillaries, 2-4 nestlings/nest, see Table 1.). When performing the brood size manipulation and cross-fostering we avoided moving the smallest or biggest nestlings to minimize sibling competition that could have significantly decreased nestlings’ survival chances after the manipulation. Body mass of nestlings swapped between nests was as similar as possible and cross-fostered individuals were kept in a warm box during the transfer (using heating pads). Nestlings were ringed 7 days after hatching, weighed and measured with a metal ruler (wing length ± 1mm) at days 7 and 14 (Table 1). Nestlings were blood sampled at day 14 (~30-75µL from the brachial vein using heparinized capillaries). Blood samples were used to (1) evaluate mitochondrial aerobic metabolism (fresh samples kept on ice collected on 14-day-old as nestlings and juveniles, Table 1), to (2) measure mitochondrial DNA copy number (i.e., mtDNA*cn)*, a proxy of mitochondrial density (measured on frozen blood samples on 2 and 14-day-old nestlings and as juveniles when samples were available), and to (3) measure mitochondrial reactive oxygen species (ROS) measured in 14-day-old nestlings and juveniles from the same samples as the mitochondrial aerobic metabolism assay (see below for detailed protocol).

Previous data on this population (Ruuskanen, *unpublished data*) showed that dispersion of great tits after fledging is almost entirely limited in this study area as none of the birds ringed as nestlings were recaptured outside of the study area. Thus, we were able to use the recapture probability of nestlings the following autumn (as juveniles, between 9 to 20 weeks after fledging) as a proxy of medium-term apparent survival. We conducted mist-nesting with playback at 6 feeding stations inside the study area (3 sessions of ca 2-4h / feeding station over October/November summing up to a total of 14 days and 69 hours of mist-nesting). Juveniles were visually sexed. In total, we recaptured 67 individuals from 34 nests: (juveniles/nests) *n_C_* = 22/9; *n_E_* = 31/15; *n_R_* = 14/10, Table 1).

### c) Mitochondrial DNA copy number

We randomly selected a minimum of 2 nestlings per nest (one original and one cross-fostered nestling). Genomic DNA was extracted from 1 to 5µL of frozen blood samples (stored at −80°C) using a salt extraction procedure adapted from Aljabani and Martinez (1997). Due to small volumes, some of the blood samples collected on day 2 could not be analysed. When data were available (see Table 1), we measured mtDNA*cn* on the same individuals at day 2, day 14 and as juvenile (i.e., recaptured in autumn 2020). DNA quantity and purity were estimated using a *NanoDrop ND-1000* spectrophotometer. Samples were re-extracted if needed ([DNA] < 50ng/µL, 260/280 ratio < 1.80 or 260/230 < 2). Samples were then diluted to 1.2ng/µL in sterile H_2_O and stored at −80°C until qPCR assays. We quantified mtDNA*cn* using real-time quantitative PCR assays (qPCR) from a protocol described in Cossin-Sevrin et al. (2022). We made some adjustments to the original protocol: samples were automatically pipetted (epMotion® 5070, Eppendorf, Hamburg, Germany) in duplicates in 384-qPCR plates (n = 5 plates) and qPCR were performed with a Biorad instrument (CFX-384, Biorad, Hercules, USA). We used Recombination Activating Gene 1 (*RAG1*) as a single control gene and cytochrome oxidase subunit 2 (*COI2*) as specific mitochondrial gene (sequences and procedure of verification are described in Cossin-Sevrin et al., 2022). qPCR reactions were conducted in a total volume of 12µL, including 6ng of DNA samples, primers at a final concentration of 300nM and 6µL of GoTaq® qPCR Mix (Promega, Madison, USA). qPCR conditions were the following : 3min at 95°C (polymerase activation), followed by 40 cycles of 10s at 95°C, 15s at 58°C, 10s at 72°C. Melting curve program was 5s at 65°C, and 0.5°C/s increased until 95°C. A pooled DNA sample from 14 adult individuals was used as a reference sample (i.e., ratio = 1.0 for mtDNA*cn*) and was included in duplicate on every plate. qPCR efficiencies of *RAG1* and *COI2* genes were respectively (mean ± SEM): 99.14 ± 1.17% and 95.74 ± 0.11%. Repeatability of mtDNA*cn* between sample-duplicates was R = 0.90 (CI 95% = [0.88, 0.92]). The samples were distributed randomly on different plates and in order to control for interplate variability, qPCR plate number was included as a random intercept in our statistical analysis (see details below). DNA integrity of 46 randomly selected samples was evaluated and deemed satisfactory using gel electrophoresis (100ng of DNA, 0.8% agarose gel at 100mV for 1 hour).

### d) Mitochondrial aerobic metabolism

In order to test the impact of brood size on nestling mitochondrial respiration, we measured mitochondrial aerobic metabolism in a subsample (1 to 3 nestlings per nest), 14 days after hatching (individuals/nest: *n_C_* = 26/14, *n_E_* = 41/21, *n_R_* = 35/19) and in the same individuals as juveniles (recaptured in autumn 2020), when samples were available (N = 14 individuals). We additionally measured mitochondrial aerobic metabolism from the majority of juveniles recaptured that participated in the manipulation (as nestlings) (in total, juvenile/nest: *n_C_* = 16/9, *n_E_* = 26/15, *n_R_* = 12/8). Blood sample volumes collected on 2-day-old nestlings were unfortunately not large enough for measuring mitochondrial aerobic metabolism at this stage (i.e., 1-10µL of blood). Mitochondrial respiration was analyzed using high-resolution respirometry (3 *Oroboros* Instruments, Innsbruck, Austria) at 40°C adapted from a protocol described in Stier et al., (2019): digitonin (20µg/mL), pyruvate (5mM), malate (2mM), ADP (1.25mM), succinate (10mM), oligomycin (2.5µM), antimycin A (2.5 µM). We used 20µL (nestlings) to 30µL (juveniles) of fresh blood when available, suspended in Mir05 buffer. Five distinct respiration rates were analysed: 1) the endogenous cellular respiration rate before permeabilization (*ROUTINE*), 2) the maximum respiration rate fueled with exogenous substrates of complex I, as well as ADP (*CI*), 3) the maximum respiration rate fueled with exogenous substrates of complexes I and II, as well as ADP (*CI+II*), 4) the respiration rate contributing to the proton leak (*LEAK*), 5) the respiration rate supporting ATP synthesis through oxidative phosphorylation (*OXPHOS*). We also calculated three mitochondrial flux ratios (FCR): 1) *OXPHOS* coupling efficiency (*OxCE = (CI+CII-LEAK)/CI+II*), 2) the proportion of maximal respiration capacity being used under endogenous cellular condition (i.e., FCR *_ROUTINE/CI+II_*) and 3) the ratio between the maximal respiration rate of complex I and the maximal respiration capacity (i.e., FCR *_CI/CI+II_*). *OXPHOS* coupling efficiency FCR provides an index of mitochondrial efficiency in producing ATP, whereas FCR *_ROUTINE/CI+II_* reflects the cellular control of mitochondrial respiration by endogenous ADP/ATP turnover and substrate availability. Respiration rates were standardized by the number of cells in each sample, measured by *BIO-RAD* TC20 automated cell counter. The technical repeatability of mitochondrial aerobic metabolism measurements was high: *ROUTINE:* R = 0.985 (CI 95% = [0.936, 0.997]); *CI+II*: R = 0.98 (CI 95% = [0.912,0.995]); *LEAK*: R = 0.979 (CI 95% = [0.916, 0.995]); *OXPHOS*: R = 0.977 (CI 95% = [0.898,0.995]) based on 9 duplicates.

### e) Reactive oxygen species measurements

Reactive oxygen species (ROS) were measured in 14-day-old nestlings and juveniles from the same samples as the mitochondrial aerobic metabolism assay (i.e., red blood cells suspended in MiR05 buffer) (see Table 1 for sample sizes). The relative amount of ROS was estimated by fluorescence, using MitoSOX™ Red kit (MitoSOX™ red mitochondrial superoxide indicator, Thermo Fisher) that specifically measures mitochondrial superoxide (i.e., the primary mitochondrial ROS) in live cells. Samples were supplemented with 4µL of MitoSOX™ (final concentration 4µM) and incubated for 30 min at 40°C protected from light. After being cooled down (5 min on ice) and centrifuged (2 min, 1000g at 4°C), samples were re-suspended in 250µL Mir05 buffer added with 5mM pyruvate, 2.5mM malate, 10mM succinate and 1.25mM ADP. 100µL of samples were loaded on a white 96-well plate (n = 43) with a transparent bottom. Kinetics of fluorescence were read for 30 min (emission 510 nm/ excitation 580 nm) in EnSpire® 2300 Multilabel Reader (PerkinElmer) set at 40°C. Samples were analyzed in duplicates. The slope of relative fluorescence (RFU/min) was then extracted and normalized by the internal control present on each plate (dry *Saccharomyces cerevisiae* diluted at 10mg/mL in Mir05). As a positive control (for mitochondrial ROS production) diluted *Saccharomyces cerevisiae* supplemented with antimycin A was included in each plate. Relative mitochondrial ROS results were standardized by the number of cells present in each well, taking into account dilution factor (cell count estimated with the *BIO-RAD* TC20 automated cell counter). Repeatability of the ROS production measurements between sample-duplicates was R = 0.924 (CI 95% = [0.9, 0.941]).

### f) Parental feeding rates

In order to test if parental feeding rates changed following the brood size manipulation, we video-recorded a subsample of nest boxes (*n_C_* = 8, *n_E_* = 15, *n_R_* = 14 nest boxes) 8 days after hatching. The cameras were concealed at ca. 2 m distance from the nest boxes. Videos were recorded for approximately 2h (mean ± SD = 137.58 ± 25.19 min) between 7 and 12 am. Standardized parental feeding rate differences (number of nest visits divided by the total length of the video starting from the first visit) was quantified using *BORIS* software (Olivier Friard & Marco Gamba, 2016), by a single observer blind to the experimental treatment.

### g) Statistical analysis

Statistical analyses were conducted using R v.4.0.2 (R core team, 2020) and performed using linear mixed models (LMMs) or general linear mixed models (GLMMs). Pre-treatment clutch sizes (raw data mean ± SEM: R = 9.24 ± 0.26, C = 8.65 ± 0.28, E = 8.48 ± 0.17 eggs; ANOVA: *F* = 2.97, *P* = 0.06) and hatching date (C = 58.70 ± 1.21, E & R = 60.16 ± 1.06 days; ANOVA: *F* = 0.54, *P* = 0.59) were relatively balanced between treatment groups. Initial brood sizes on day 2 post-hatching per treatment groups were the following: (raw data mean ± SEM [range]) R = 8.00 ± 0.32 [5;11] chicks, C = 7.50 ± 0.44 [4;10] chicks and E = 7.68 ± 0.28 [4;9] chicks and were not statistically different between treatment groups before the manipulation (ANOVA: *F* = 0.55, *P* = 0.57).

#### Experimental approach

To investigate the experimental effect of brood size manipulation on response variables (i.e., body mass, wing length, mtDNA*cn*, mitochondrial aerobic metabolism, mitochondrial ROS production), we always included in our models the treatment as a 3-level fixed factor (R,C,E) and the initial brood size as a continuous variable to account for initial differences in brood size across nests. These analyses are referred to “*experimental approach”* in the text. To test for potential different effects of the treatment according to the initial number of nestlings in the nest, we always tested the interaction between the treatment and initial brood size in our models. Non-significant interaction (treatment* initial brood size) and cross-fostering status (i.e., cross-fostered or not, included as main effect in models) were dropped (starting from the interaction) from the models in a backward-stepwise procedure to obtain the lowest Akaike Information Criterion (AIC) value. When AIC were similar between models (differences between AIC less than 2), we chose the simplest model (with the lowest degree of freedom). For models that included repeated measures across time (i.e., see below body mass), we initially included the age, treatment, initial brood size and their interaction and removed non-significant interactions following a backward-stepwise procedure. For changes in mt*DNAcn* with time (from day 2 to 14), we present results from the treatment and age interaction (although non-significant), as we predicted an effect of the treatment with time. However, the initial brood size could not be included as a fixed factor in the model because of convergence issues. We also included bird ID as a random intercept to take into account the non-independence of measures from the same individual. Unfortunately, only a few nestlings measured at day 14 for mitochondrial respiration rates were recaptured as juveniles, thus we could not add the bird ID as a random intercept for mitochondrial respiration traits in our models (convergence issues).

#### Correlative approach

To explore the associations between number of nestlings and the measured traits (focusing on the ecological aspect of the brood size rather than experimental), we used another set of models including the actual number of nestlings (on the day of data collection) as a continuous variable. These analyses are referred to “*correlative approach”* in the text. As the number of nestlings per nest nests varied substantially across and within treatment groups (e.g., at day 14 brood size ranged from 2 to 11 nestlings), this analysis reflects the associations between a given brood size and trait of interest. However, given that the dataset using brood size as a continuous variable includes both experimentally manipulated (E, R) and non-manipulated nests (C) we also analyzed the associations between the number of nestlings and target variables using only the non-manipulated nests (C) group to check if patterns might have been confounded by including experimental nests (see ESM.A). As results were similar (ESM.B Table 2), we report results of the full dataset in the main text. In both analyses, we included hatching date as a continuous variable and the IDs of both original and rearing nest boxes as random intercepts. qPCR plate ID could not be included in the model only including the control group because of convergence issues.

**Table 2.** Results of a LMM testing the effect of age and brood size manipulation treatment on nestling body mass. Day 2: n = 540 observations, day 7: n = 420 observations, day 14: n = 403 observations, N = 540 individuals in total. Estimates are reported with their 95% CI. Chick ID (ring), original nest box ID and nest box of rearing ID were included as random intercepts in models. σ2, within-group variance; τ00, between-group variance. Sample size (n) along with marginal (fixed effects only) and conditional (fixed and random effects). Bold indicates significance (P < 0.05).

| Predictor | Estimate | 95% CI | P value |
| --- | --- | --- | --- |
| (Intercept) | -0.27 | -2.37 – 1.83 | 0.799 |
| age (day 7) | 7.69 | 7.38 – 7.99 | **<0.001** |
| age (day 14) | 13.67 | 13.36 – 13.98 | **<0.001** |
| treatment (E) | -0.08 | -0.54 – 0.39 | 0.748 |
| treatment (R) | -0.11 | -0.58 – 0.37 | 0.659 |
| hatching date | 0.05 | 0.02 – 0.09 | **0.003** |
| age (day 7) : treatment (E) | -0.21 | -0.58 – 0.17 | 0.288 |
| age (day 14) : treatment (E) | 0.05 | -0.34 – 0.43 | 0.806 |
| age (day 7) : treatment (R) | 0.12 | -0.30 – 0.54 | 0.577 |
| age (day 14) : treatment (R) | 0.78 | 0.35 – 1.20 | **<0.001** |
| Random effects |  |  |  |
| $\sigma^2$ | 1.35 | | |
| $\tau_{00}$ ring | 0.13 | | |
| $\tau_{00}$ nest of origin | 0.36 | | |
| $\tau_{00}$ nest of rearing | 0.13 | | |
| n ring | 540 |  |  |
| n nest of origin | 70 |  |  |
| n nest of rearing | 70 |  |  |
| n observations | 1362 |  |  |
| Marginal R<sup>2</sup> / Conditional R<sup>2</sup> | 0.945/0.962 |  |  |

Standardized parental feeding rate differences were tested according to treatment groups and the initial brood size, but also according to the number of nestlings at day 7, using in both cases a linear model without random effects (LM). We included the starting time of the video recordings as a covariate in models to account for differences in feeding rates during the day.

Nestling growth metrics (i.e., postnatal body mass and wing length) were analyzed using LMMs with both the original nest box ID and the nest box of rearing ID as random intercepts. For longitudinal measurements, we included bird ID as a random intercept.

mtDNA*cn* data distribution did not fulfill the criteria of normality according to a Cullen and Frey plot (*fitdistrplus* package; Delignette-Muller and Dutang, 2015); therefore, we analyzed the effects of the treatment and the number of nestlings on mtDNA*cn* using a GLMM (gamma error distribution, log link). We included the qPCR plate ID as a random intercept. For juveniles, we tested the association between mtDNA*cn* and the number of nestlings in the nest a few days before fledging, by adding the brood size at day 14 as explanatory factor in our model (GLM, gamma error distribution, log link). All mitochondrial respiration rates (recorded on 14-day-old nestlings and juveniles, including *ROUTINE, CI, CI+II, LEAK, OXPHOS*) were tested with LMMs. We analyzed mitochondrial respiration rates at the mitochondrial level (i.e., respiration measurements controlled for mitochondrial density by inclusion of mtDNA*cn* as a covariate), which indicates the respiration rate per unit of mitochondria. For mitochondrial respiration rates measured at day 14, we further quantified the variance explained by the random intercepts (i.e., both original nest box ID and nest box of rearing ID included as random intercepts, while treatment, initial brood size, hatching date and mtDNA*cn* were included as fixed factors), using *RptR* package (gaussian distribution, N bootstraps = 1000) (Nakagawa & Schielzeth, 2010; Stoffel et al, 2017). Mitochondrial ROS production in nestlings (day 14) and juveniles was analyzed according to the treatment and the initial brood size, but also according to the number of nestlings at day 14 using a LMM.

The effect of the brood size manipulation and the number of nestlings on survival metrics (fledging success and recapture probability as juveniles) were estimated with GLMMs (logistic binary distribution of dependent variables: 0 = dead, 1 = alive). We included hatching date as covariate, while both original nest box ID and nest box of rearing ID were included as random intercepts. In case of convergence issues with the models, we only included the nest of rearing ID as a random intercept and removed the hatching date from covariates if needed.

For investigating the contribution of mitochondrial respiration rates at day 14 on juvenile apparent survival (i.e., recapture probability), we performed GLM on survival (logistic binary distribution of dependent variables: 0 = dead, 1 = alive) and included mitochondrial respiration rates or FCR(s) and hatching date as explanatory factors. As the number of individuals recaptured was less than 2 individuals for several nests, we could not include the nest of rearing ID as a random intercept in our models (convergence issues).

All models were performed using *lme4 package* (Bates et al., 2015). Results from type III ANOVA tables with *F* values and *P* values (i.e., testing the main effect of each factor and interaction) were calculated based on Satterthwaite’s method and are presented in the text. Results from GLMMs (logistic binary distribution) were calculated based on Wald Chisquare tests (type II ANOVA). Model estimates and Odds Ratios (with associated 95% CI and *P* values) are reported in tables. *emmeans* package was used for conducting multiple *post hoc* comparisons (adjusted with Tukey honest significant differences correction). Effect-sizes (Cohen’s D) were estimated using *effsize* package (Ben-Shachar et al., 2020). Values were considered as statistically significant for P < 0.05.

## Results

### 1. Brood size manipulation

Our treatment led to significant differences in brood size between treatment groups (R, C, E) after the manipulation: average (± SEM, on raw data) brood sizes were R = 6.00 ± 0.32 (initial 8.00 ± 0.32), C = 7.50 ± 0.44 (initial 7.50 ± 0.44), E = 9.68 ± 0.28 (initial 7.68 ± 0.28) nestlings per nest on day 2 (Tukey HSD *post hoc:* all comparisons *P* < 0.009). Brood size remained significantly higher for the E group than C or R during the whole growth period (from day 2 to day 14) (all Cohen’s D > 1.50) (Tukey HSD *post hoc:* C vs. E and E vs. R comparisons, all *P* < 0.02), while the differences in brood sizes between C and R groups were not significant at 7 days (Cohen’s D with 95% CI = 0.43 [-0.25, 1.11]) and 14 days after hatching (Cohen’s D with 95% CI = 0.37 [-0.31, 1.05]) (Tukey HSD *post hoc:* C vs. R comparison, all *P* > 0.90). Averages (± SEM, on raw data) for R, C and E groups were respectively: R = 4.84 ± 0.54, C = 5.25 ± 0.72, E = 7.88 ± 0.76 nestlings at day 7 and R = 4.60 ± 0.54, C = 4.95 ± 0.68, E = 7.56 ± 0.75 nestlings at day 14.

### 2. Parental feeding rates and nestling growth trajectories

#### 2.1. Experimental approach

Parental feeding rate (8 days after hatching) was significantly affected by the treatment (*F_2, 32_* = 4.64, *P* = 0.02, see Fig.2A) with higher rates for the E group (raw data mean ± SE = 41.26 ± 6.03 visits per hour) compared to R group (raw data mean ± SE = 25.75 ± 4.05) (Tukey HSD *post hoc* comparison: *P* = 0.04). Differences in parental feeding rate between E and C groups (C: raw data mean ± SE = 28.49 ± 5.22) were close to significance (Tukey HSD *post hoc* comparison: *P* = 0.051). Parental feeding rate significantly increased with initial brood size (estimate ± SE = 2.76 ± 1.55, *F_1,32_* = 7.91, *P* = 0.008) and significantly decreased with time of day (estimate ± SE = −2.67 ± 6.13e-10, *F_1,32_* = 19.01, *P* < 0.001).

**Fig. 2:**
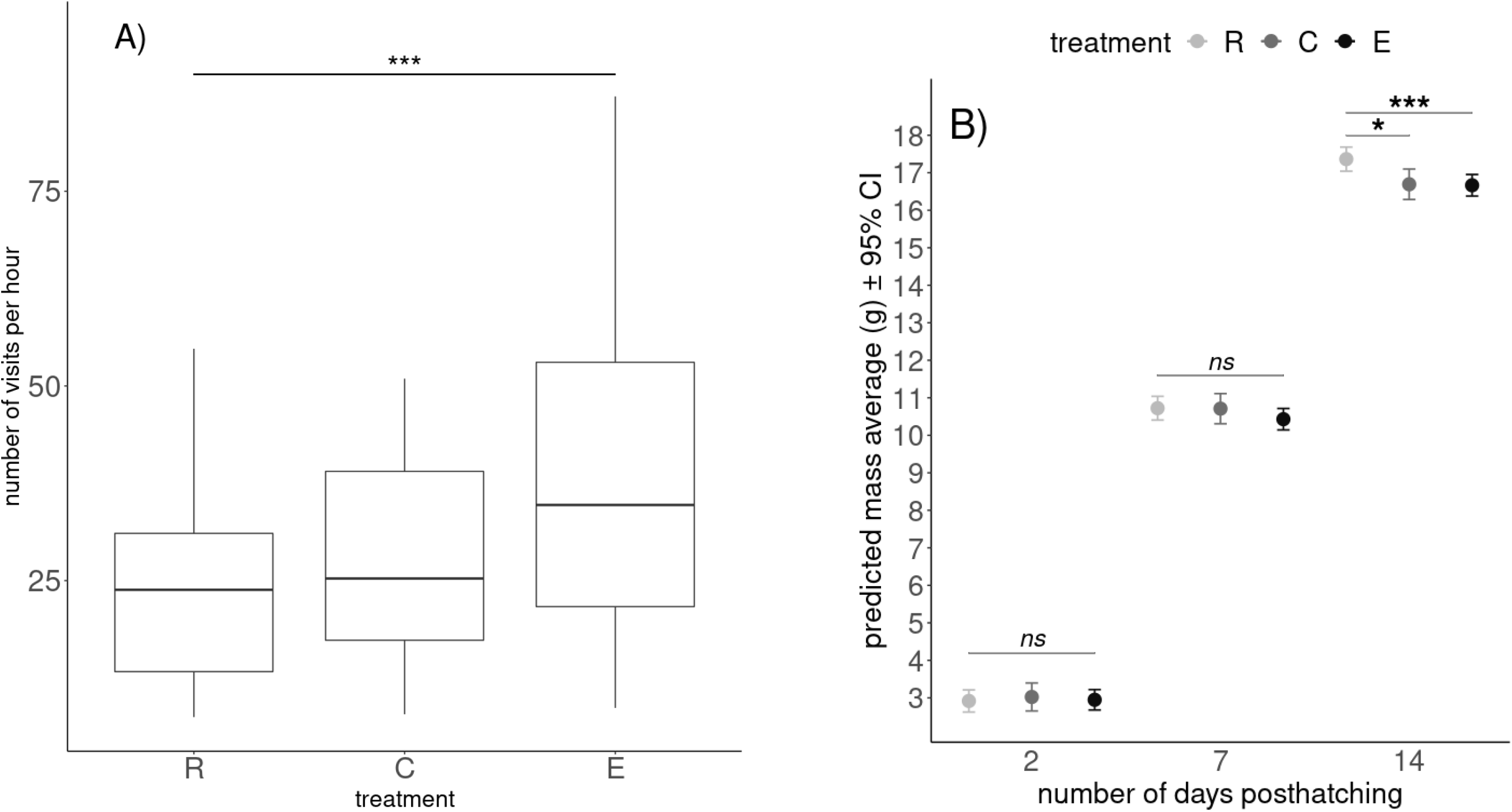
Parental feeding rate (A) and predicted body mass average of nestlings during the growth period (B) according to brood size manipulation treatment groups: reduced (R), control (C), enlarged (E) brood sizes. For A), raw data distribution is presented with boxplots (*n_C_* = 8, *n_E_* = 15, *n_R_* = 14 nest boxes). Stars indicate the significance of Tukey HSD *post hoc* test (*** *P* < 0.001). R2 = 0.53. For B), predicted values with their 95% CI and results from Tukey HSD *post hoc* tests are reported. Stars indicate the significance of the *post hoc* test (*** *P* < 0.001, * *P* < 0.05). R2 = 0.96. See Table 1 for sample-sizes.

Postnatal body mass dynamic (from day 2 to 14) was differentially affected by the treatment depending on offspring age (day 2, day 7 and day 14: age*treatment: *F_4,930.28_* = 5.07, *P* < 0.001, Table 2). Specifically, nestlings from the R group had a higher body mass 14 days after hatching than nestlings from C (+3.86%) and E groups (+3.97%) (Tukey HSD *post hoc* R vs. C and R vs. E comparisons: all *t* < −2.55, all *P <* 0.03, see Fig. 2B), while body mass at day 14 from nestlings raised in C and E groups were similar (Tukey HSD *post hoc* C vs. E comparison: t = 0.11, *P* = 0.99, see Fig.2B). We did not find any significant difference in body mass 2 and 7 days after hatching (Tukey HSD *post hoc* comparisons: all *t* < 1.12, all *P >* 0.50). Body mass significantly increased with hatching date (*F_1,79.19_* = 9.61, *P* = 0.003, see Table 2). The treatment did not significantly impact nestling wing length during the growth period (day 7 and day 14) (all *F* < 0.68, all *P >* 0.51). We found a significant positive correlation of wing length and initial brood size at day 14 (estimate ± SE = 0.42 ± 0.18, *F_1,41.5_* = 5.66, *P* = 0.02). At both ages (day 7 and 14), wing length significantly increased with hatching date (all *F* > 6.57, all *P* < 0.01). Juvenile body mass and size were not associated with the treatment, the initial brood size nor both in interaction (all *F* < 0.62, all *P* > 0.55).

#### 2.2 Correlative approach

Parental feeding rate significantly increased with the number of nestlings recorded 7 days after hatching (estimate ± SE = 4.28 ± 1.01, *F_1, 34_* = 22.41, *P* < 0.001).

When analyzing each age separately, in order to account for the number of nestlings in the nest at a given age, nestling body mass at day 7 was negatively associated with the number of nestlings in the nest (estimate ± SE = −0.16 ± 0.06, *F_1, 45.44_* = 6.15, *P* = 0.02), while we did not find an association for the wing length (*F_1, 31.10_* = 0.38, *P* = 0.54). At day 14, nestling body mass was not significantly associated with the number of nestlings (*F_1, 52.70_* = 0.12, *P* = 0.73). Nestling wing length at day 14 tended to increase with the number of nestlings (estimate ± SE: 0.23 ± 0.11, *F_1, 35.58_* = 4.02, *P* = 0.05, see Fig.3). Nestling body mass and wing length both significantly increased with the hatching date at day 7 and 14 (all *F* > 5.12, all *P* < 0.03).

**Fig. 3.**
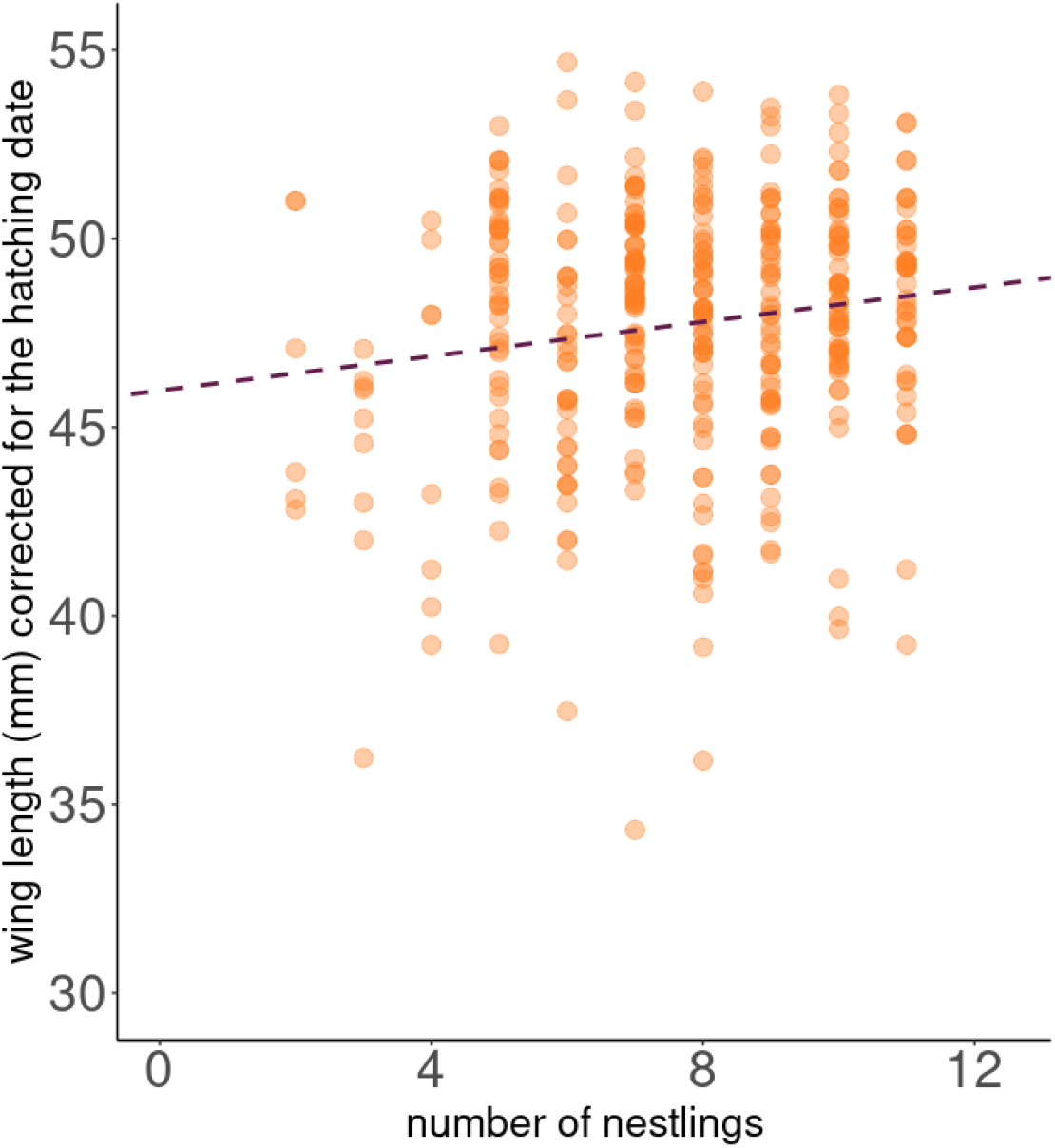
Predicted values of the wing length of 14-days-old nestlings according to the number of nestlings in the nest at day 14. Predicted values are extracted from linear mixed models (LMMs) and corrected for the average hatching date of the season. Regression line (in dotted line) and results from the models are presented. N = 403 individuals. Conditional R2 of the model presented was 0.65.

### 3. Mitochondrial DNA copy number

#### 3.1. Experimental approach

While mtDNA*cn* was not significantly impacted by the interaction of the age and the treatment (χ2 = 0.03, *P* = 0.11), mtDNA*cn* significantly decreased during the entire growth period (from day 2 to 14: Cohen’s D with 95% CI = 1.88 [1.54, 2.21]) (estimate ± SE = −0.1 ± 0.01, *P* < 0.001, juveniles not included in the repeated measures analysis because of limited sample size). Juvenile mtDNA*cn* was not significantly impacted by the treatment or the initial brood size (all *P* > 0.6).

#### 3.2. Correlative approach

While mtDNA*cn* at day 14 was not associated with the number of nestlings in the nest (*P* = 0.11), larger brood sizes a few days before fledging (i.e., day 14) predicted higher mtDNA*cn* for juveniles (estimate ± SE = 0.07 ± 0.03, *P* = 0.04).

### 4. Mitochondrial aerobic metabolism

#### 4.1. Experimental approach

We did not find any significant effect of the brood size manipulation treatment or of the initial brood size on the different mitochondrial respiration rates and FCR(s) measured at day 14 (all *F <* 2.17, all *P* > 0.13, Fig.4). Juvenile mitochondrial respiration rates and FCR(s) were not significantly impacted either by the treatment (all *F <* 0.75, all *P* > 0.48) or the initial brood size (all *F* < 2.36, all *P >* 0.13). All mitochondrial respiration rates increased with mtDNA*cn* at day 14 (all *F >* 65.14, all *P <* 0.001) and in juveniles (all *F* > 5.39, all *P >* 0.02), except for *LEAK* (juveniles: *F_1, 49_* = 3.07, *P* = 0.09).

**Fig. 4:**
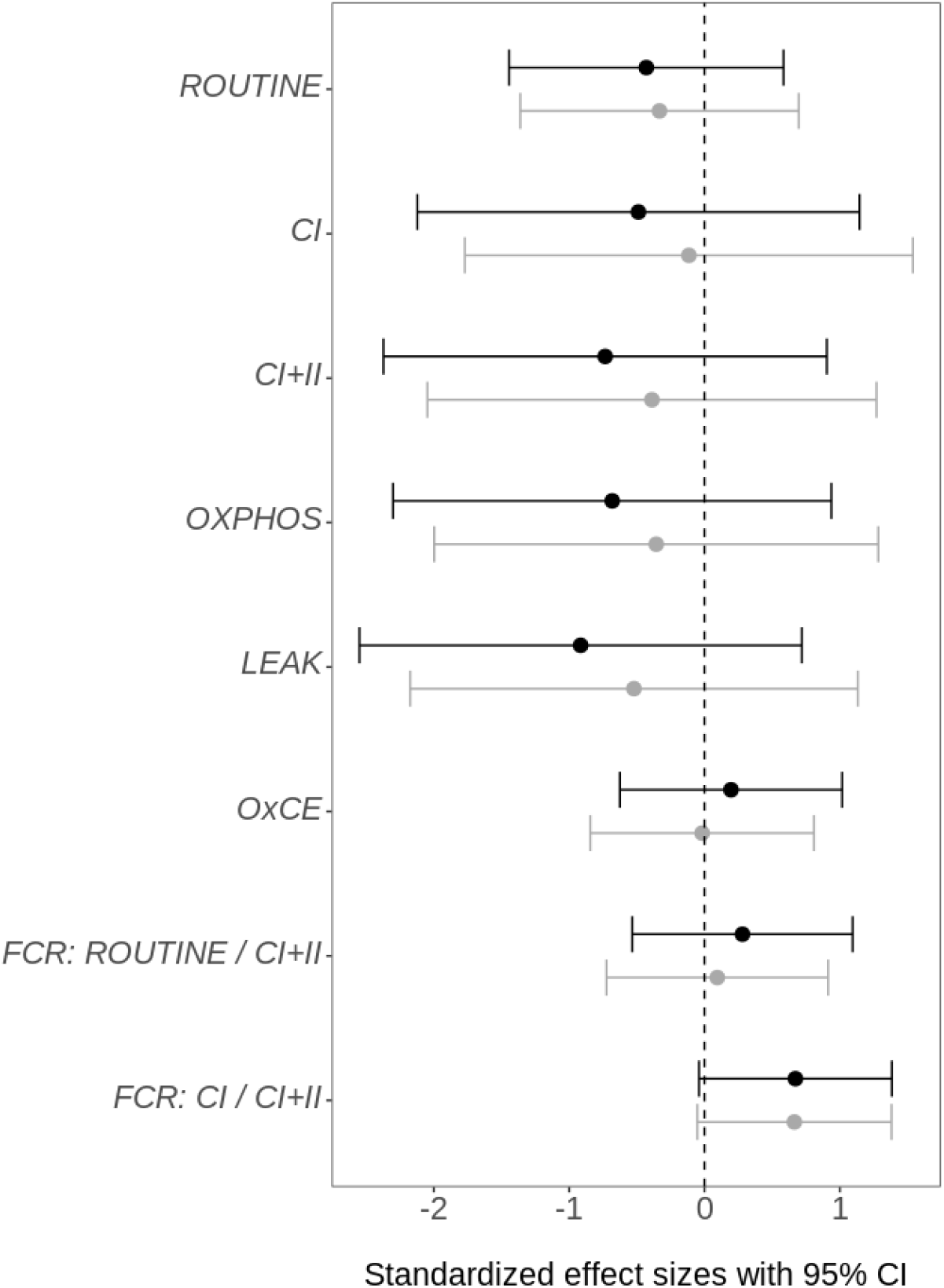
Effect of the brood size manipulation on mitochondrial metabolic rates and flux control ratios. Mitochondrial aerobic metabolism was measured at day 14 between individuals raised in reduced, control and enlarged broods (see sample-sizes Table 1). Standardized effect sizes are based on predicted values of the model and reported with their 95% CI. In black, effect sizes between individuals raised in enlarged vs. control broods. In grey, effect sizes between individuals raised in reduced vs. control broods.

For all mitochondrial respiration rates measured at day 14, the nest of rearing significantly contributed to explain the variance in our models (all repeatabilities > 0.51, all *P* < 0.001, see Fig.5). Except for *ROUTINE* (repeatability = 0.08, *P* = 0.20), the variance explained by the nest of origin was significantly higher than 0 (all repeatabilities > 0.13, all *P* < 0.02) but the contribution of the nest of rearing was higher than the nest of origin (Fig.5).

**Fig. 5:**
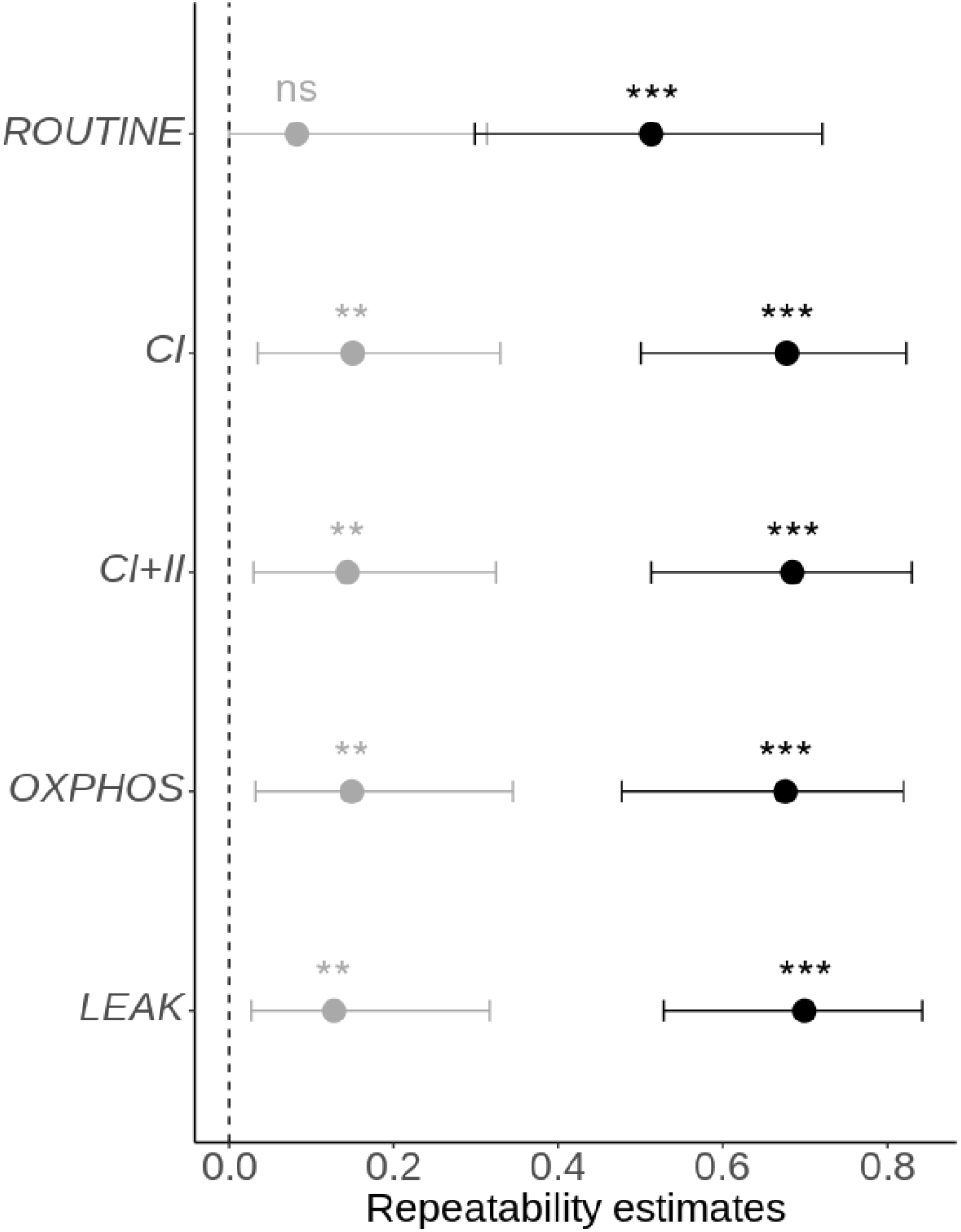
Variance explained by the nest of origin (in grey) and the nest of rearing (in black) in linear mixed models testing mitochondrial respiration rates at day 14 according to the number of nestlings (at day 14). Stars indicate significance to be different from 0 (*** *P* < 0.001, ** *P* < 0.01). Repeatabilities are presented with their 95% CI. ns: non significant. See Table 1 for sample-sizes.

#### 4.2. Correlative approach

We found a negative association between the number of nestlings at day 14 and mitochondrial respiration rates measured at day 14 (all *F* > 8.80, all *P* < 0.005, see Table 3, Fig.6). *OXPHOS* coupling efficiency and both FCR *_ROUTINE/CI+II_* and FCR *_CI/CI+II_* were not significantly associated with the number of nestlings at day 14 (all *F* < 1.37 and all *P* > 0.25, see ESM.A). We found similar results when only including individuals raised in the C group (see ESM.B, Table 2). *CI, CI+II, OXPHOS* and *OXPHOS* coupling efficiency all significantly decreased with the hatching date (all *F >* 9.58, all *P <* 0.003). *ROUTINE, CI, CI+II, LEAK* and *OXPHOS* significantly increased with *mtDNAcn* (all *F >* 63.49, all *P* < 0.001, see Table 3). Since nestlings from very small brood sizes had higher mitochondrial respiration rates (see Fig.6), which could drive the associations, we performed the same statistical analysis excluding nestlings raised in small broods (less than 5 chicks 14 days post hatching) (n = 28 nestlings from 12 nests removed from the analysis). In this case, we could not detect any significant associations between the number of nestlings (day 14) on the different mitochondrial respiration rates measured (all *F <* 2.23, all *P* > 0.14, see ESM.B). Juvenile mitochondrial respiration rates (all *F <* 0.21, all *P* > 0.65) or FCRs (all *F <* 0.72, all *P >* 0.49), were not associated with the number of nestlings at day 14, except for FCR *_CI/CI+II_* for which we found a negative association (estimate ± SE = −0.005 ± 0.003, *F_1, 62_* = 4.36, *P* = 0.04). *ROUTINE, CI, CI+II* and *OXPHOS* significantly increased with juvenile mtDNA*cn* (all *F* > 5.26, all *P* < 0.03), while *LEAK* was not significantly associated with mtDNA*cn* (*F_1, 51_* = 1.95, *P* = 0.17).

**Fig. 6.**
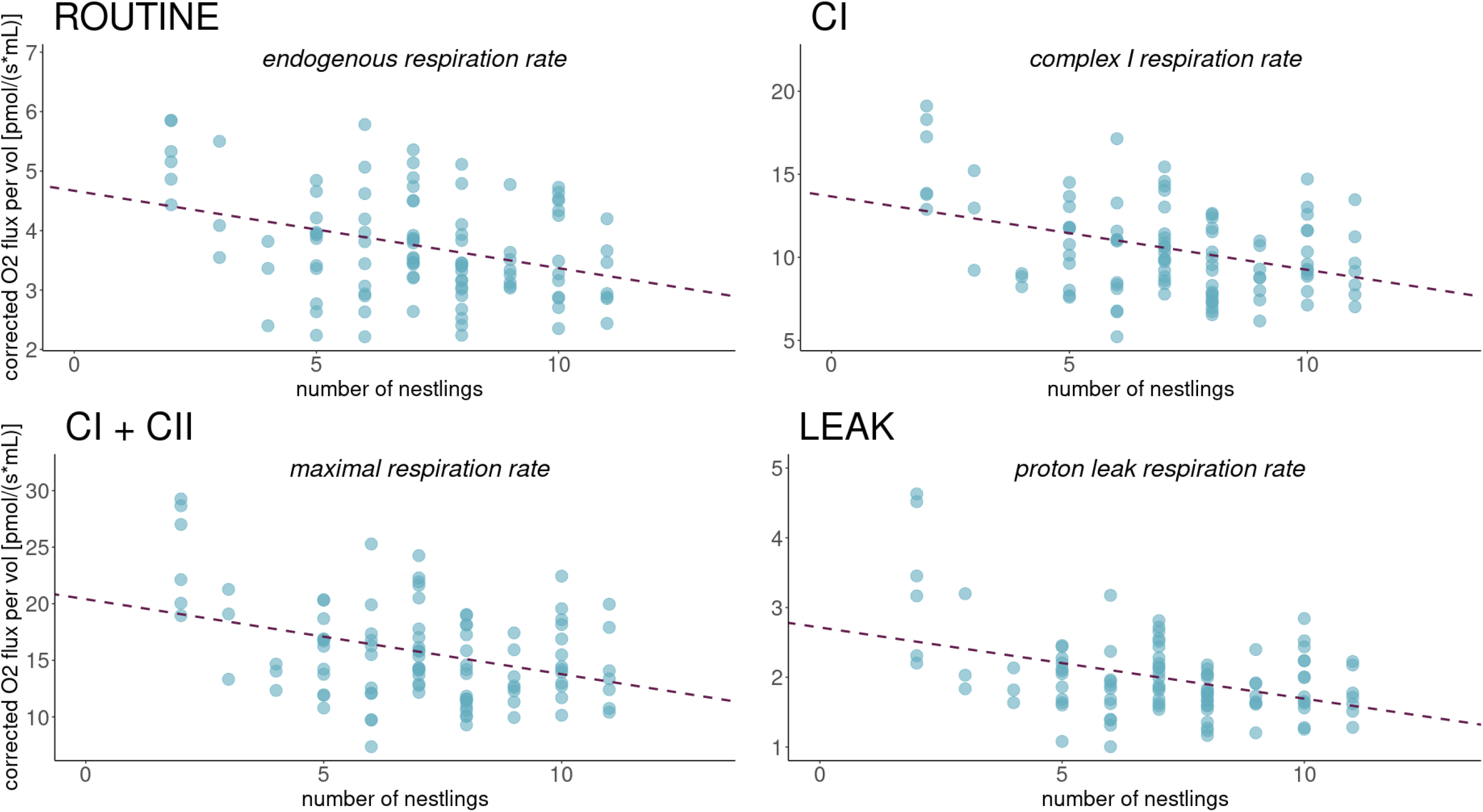
Predicted values of mitochondrial respiration rates on 14 days old nestlings according to the number of nestlings at day 14. N = 102 individuals. Predicted values are extracted from linear mixed models (LMMs). Regression lines (in dotted lines) and results from the models are presented. Predicted values are corrected for the average hatching date of the season. Mitochondrial respiration rates were corrected for mitochondrial DNA copy number (i.e., proxy of the mitochondrial density). Original nest box ID and nest box of rearing ID were included as random intercepts in the models. R2 of each model are reported in Table 3.

**Table 3.** Results of linear mixed model testing the associations between the number of nestlings in the nest (14 days after hatching) and mitochondrial respiration rates measured on 14-day-old nestlings (N = 102 individuals, n = 55 nest boxes). Mitochondrial respiration rates were corrected for the mitochondrial DNA copy number (i.e., proxy of mitochondrial density). Linear mixed models (LMM) estimates are reported with their 95% CI. Original nest box ID and nest box of rearing ID were included as random intercepts in the models. σ2, within group variance; τ00 between-group variance. Bold indicates significance (*P* < 0.05).

|  | ROUTINE |  |  | CI |  |  | CI + II |  |  | LEAK |  |  |
| --- | --- | --- | --- | --- | --- | --- | --- | --- | --- | --- | --- | --- |
| Predictors | Estimates | CI 95% | P-value | Estimates | CI 95% | P-value | Estimate<br>s | CI 95% | P-value | Estimates | CI 95% | P-value |
| (Intercept) | 4.55 | 2.37 – 6.72 | **<0.001** | 20.12 | 12.93 - 27.31 | **< 0.001** | 29.39 | 18.21 – 40.57 | **<0.001** | 2.70 | 1.20 – 4.20 | **<0.001** |
| number of nestlings | -0.13 | -0.22 – -0.04 | **0.005** | -0.44 | -0.72 – -0.17 | **0.002** | -0.66 | -1.09 – -0.23 | **0.003** | -0.10 | -0.16 – -0.04 | **<0.001** |
| mtDNA$_{cn}$ | 0.34 | 0.25 – 0.42 | **<0.001** | 0.91 | 0.69 – 1.12 | **<0.001** | 1.44 | 1.10 – 1.77 | **<0.001** | 0.18 | 0.14 – 0.23 | **<0.001** |
| hatching date | -0.02 | -0.06 – 0.02 | 0.305 | -0.17 | -0.29 – -0.04 | **0.009** | -0.24 | -0.43 – -0.05 | **0.013** | -0.01 | -0.04 – 0.01 | 0.384 |
| Random effects |  |  |  |  |  |  |  |  |  |  |  |  |
| $\sigma^2$ | 0.32 | | | 1.31 | | | 3.13 | | | 0.06 | | |
| $\tau_{00}$ nest of origin | 0.05 | | | 1.10 | | | 2.52 | | | 0.04 | | |
| $\tau_{00}$ nest of rearing | 0.33 | | | 4.21 | | | 10.35 | | | 0.19 | | |
| Observations | 102 |  |  | 102 |  |  | 102 |  |  | 102 |  |  |
| Marginal R<sup>2</sup> / Conditional R<sup>2</sup> | 0.488 / 0.767 |  |  | 0.487 / 0.898 |  |  | 0.483 / 0.899 |  |  | 0.454 / 0.889 |  |  |

### 5. ROS production

#### 5.1. Experimental approach

In 14-days-old nestlings, mitochondrial ROS production was not significantly affected by the treatment (*F_2, 45.7_* = 0.62, *P* = 0.54, see ESM.D) or the initial brood size (*F_1, 49.9_* = 0.05, *P* = 0.82, see ESM.D). These results remained consistent in juveniles (treatment: *F_2, 48_* = 1.58, *P* = 0.22; initial brood size: *F_1, 48_* = 0.74, *P* = 0.39, see ESM.D). While mitochondrial ROS production was not significantly associated with mtDNA*cn* in nestlings (*F_1, 83_* = 0.48, *P* = 0.49), juvenile mitochondrial ROS production significantly increased with mtDNA*cn* measured in autumn (estimate ± SE = 0.003 ± 0.001, *F_1, 48_* = 4.60, *P* = 0.04).

#### 5.2. Correlative approach

We did not find significant associations between the number of nestlings at day 14 and nestling mitochondrial ROS production (day 14: *F_1, 53.49_* = 0.42, *P* = 0.52) or in juveniles (*F_1, 50_* = 1.08, *P* = 0.30).

### 6. Survival metrics

#### 6.1. Experimental approach

Fledgling success was not significantly affected by the treatment (χ2 = 3.20, *P* = 0.25, raw data: R = 75.33%, C = 65,79%, E = 77.78%), neither by the initial brood size (χ2 = 0.006, *P* = 0.83) or the hatching date (χ2 = 2.11, *P* = 0.13). Juvenile recapture probability was not significantly affected by the treatment (χ2 = 2.27, *P* = 0.33, raw data: R = 12.17%, C = 22.22%, E = 18.52%) or the initial brood size (χ2 = 0.02, *P* = 0.87), but was negatively associated with the hatching date (χ2 *=* 15.47, *P* < 0.001).

#### 6.2. Correlative approach

Fledgling success was strongly positively associated with the number of nestlings in the nest at day 14 (χ2 = 61.47, *P <* 0.001). Juvenile recapture probability was not significantly associated with the number of nestlings day 14 (χ2 = 0.23, *P* = 0.63).

Finally, we did not find any significant associations between juvenile recapture probability, mitochondrial respiration rates and FCR(s) measured at day 14 (all *P* > 0.2, see ESM.E).

## Discussion

Overall, the experimental brood size manipulation did not significantly affect nestling mitochondrial density, metabolism or ROS production. Despite a mild impact of the treatment on nestling growth trajectories, body mass differences cannot be associated here with variation in mitochondrial metabolism. Furthermore, we did not detect any significant long-lasting effect of the brood size manipulation treatment on juveniles (neither on recapture probability, body mass and size, nor mitochondrial density, metabolism and subsequent ROS production). However, our results emphasized the importance of chick numbers in the nest regardless of experimental manipulation for nestling mitochondrial respiration. Nestling mitochondrial metabolic rates were negatively associated with the number of nestlings in the nest (but see precautions in interpretations below). Our results also provide evidence that environmental conditions during the growth period (nest of rearing) contribute more to explaining variance in red blood cells mitochondrial metabolism than genetic inheritance pre- and early postnatal parental effects (nest of origin) in great tits. Taken together, our results suggest that the actual number of nestlings (rather than the modification of initial brood size) is an important influence on nestling growth pattern and mitochondrial metabolism. The number of siblings in a nest is expected to influence food availability and competition between chicks, as well as early-life conditions critical to nestling growth, such as nest temperature (Andreasson et al., 2016; Hope et al., 2021; Nord & Nilsson, 2011).

### Experimental approach

Nestling growth trajectories (postnatal body mass) differed according to nestling age and our treatment. As expected, individuals raised in the R group had a higher body mass a few days before fledging compared to other groups (see also Hõrak, 2003). While we expected nestlings raised in E group to have lower body mass (Hõrak, 2003; Rytkönen & Orell, 2001; Smith et al., 1989), nestlings raised in E and C groups had similar body masses over the entire growth period. Moreover, nestling wing length did not differ between treatment groups. It is possible that parents managed to compensate for the brood size augmentation by increasing parental effort, as suggested by results on parental feeding rates (measured on a subsample of nests). The number of visits was significantly higher in E group compared to R and tended to be higher compared to C (although non-significant). These results would be supported by prior studies suggesting that parents can rear more nestlings than the number of eggs laid (Casti, 2018; Monaghan & Nager, 1997; Vander Werf, 1992).

It is worth noting that in our experiment the difference in nestling number between C and R groups did not remain significant (small effect-sizes between groups) at the end of the growth period (from day 7 to 14). This likely contributes to explain why our experiment failed to demonstrate large differences between treatment groups. It is interesting that even without differences in the number of chicks at the end of the experiment between C and R groups, the R group had larger chicks (see hypothesis below).

It has been shown that a brood size enlargement can affect nestling metabolism, as brood size decreases whole animal resting rate of oxygen consumption in the short-term (tree swallow), and increases standard metabolic rate in the a long-term (zebra finches) (Burness et al., 2000; Verhulst et al., 2006). In our case, the brood size manipulation treatment did not have an effect on nestling red blood cell mitochondrial metabolism during the growth period or in a longer-term in juveniles. This lack of effects may be explained by the two reasons mentioned above (i.e., increase of parental feeding rates and no differences in chick number between C and R groups). Nestling ROS production (and juvenile ROS production) were not either impacted by the treatment. This outcome is in accordance with our findings that mitochondrial aerobic metabolism did not differ between treatment groups. Despite the mild effect of brood size manipulation on nestling body mass, nestling fledgling success and apparent medium-term survival (i.e., recapture probability as juvenile) were not significantly impacted by the treatment.

### Correlative approach

Whereas the brood size manipulation treatment had only a mild effect on nestling growth pattern, our results suggest that the actual number of offspring in the nest has an important influence on nestling postnatal body mass and structural size. Nestling body mass was negatively associated with the number of nestlings in the nest in the middle of the growth period (day 7), while nestling wing length tended to be positively associated with the number of individuals in the nest at the end of the growth period (day 14). This insight was surprising as the opposite results (i.e., negative association between the wing length and the number of chicks in the nest) have been reported in the literature (Hõrak, 2003; Rytkönen & Orell, 2001; Smith et al., 1989). Yet, these results from previous studies have been found in the framework of a brood size manipulation and did not strictly focus on the actual number of chicks in the nest.

We found a negative association between mitochondrial metabolism (*ROUTINE, CI, CI+II, LEAK* and *OXPHOS)* and number of nestlings. As both *LEAK* and *OXPHOS* were negatively correlated with number of nestlings, we did not find an association between *OXPHOS* coupling efficiency and nestling number. This suggests that higher mitochondrial metabolic rates for nestlings raised in small broods were linked to an increase in oxidative phosphorylation (i.e., a proxy of ATP production) that may reflect higher energetic demands compared to larger nests. While we cannot here strictly test what requires higher energetic demands for the nestlings, it is possible that higher mitochondrial metabolic rates were linked to a higher thermogenesis associated with the small number of chicks in the nest (Bicudo et al., 2001).

While these results are in accordance with our predictions (decrease in mitochondrial metabolic rates in larger broods) it is important to note that these negative associations (nestling structural size and mitochondrial metabolism) with the number of nestlings did not remain significant when nestlings from small broods (less than 5 nestlings at day 14) were excluded from the analysis, meaning that those specific broods drove the patterns. Lower mitochondrial metabolic rates in larger broods were probably not associated with a stressful rearing environment in our case. Interestingly, broods with less than 5 nestlings at day 14 (n = 20 nests) had really low survival chances during the growth period (from day 2 to 14) compared to the larger broods (> 4 nestlings, n = 50 nests) (average on raw data: 25.5% vs. 92.4% of survival at day 14) and most of the nestlings did not reach day 7 (average at day 7: 5.1 nestlings lost in small broods vs. 0.34 in larger broods). We therefore suspect nestling growth and mitochondrial metabolic patterns to rather reflect unusual rearing conditions than being general patterns. Several hypotheses could explain higher mitochondrial metabolic rates for individuals raised in (very) small broods. Our main hypothesis is that these individuals might be at a less-advanced developmental stage. It has been shown in several avian species that mitochondrial quantity and/or respiration decreases during postnatal development (Stier et al. 2020; Stier et al. 2022; Cossin-Sevrin et al. 2022, Hsu et al. 2023; but see: Dawson & Salmón, 2020), and it is thus possible that higher metabolic rates in very small broods reflect that their nestlings are less developed for a given age. This hypothesis is supported by the fact that individuals raised in small broods had a smaller structural size (wing length) than in larger broods.

Then, the high nestling mortality may be an indication of poor rearing conditions (e.g., food quality, incubation time). It has been previously shown that in some cases environmental stressors may lead to higher metabolic rate (in interaction with glucocorticoid levels in zebra finches) (Jimeno et al., 2017).

Finally, these small broods with a high unusual mortality during early-growth may be subject to selective disappearance and nestlings surviving until 14 days after hatching represent a non-random pool of individuals that managed to survive and cope with detrimental conditions during early-growth. This hypothesis would be supported by our results showing that early-life environmental conditions are the major determinant in nestling mitochondrial metabolism in red blood cells. Indeed, our study demonstrates that both genetic inheritance (but also complementary mechanisms, such as parental effects before the cross-fostering) and the rearing environment contribute to variation in offspring mitochondrial traits, but with a larger contribution from the rearing environment. Similar results about lower contribution of familial background have been found for resting metabolic rate in collared flycatcher nestlings (*Ficedula albicollis*) (McFarlane et al., 2021). While the underlying mechanisms of modulation of mitochondria by early-life environmental conditions are unknown, recent research points out that mitochondrial function can respond to environmental cues through changes in gene expression and mitochondrial DNA methylation (Sharma et al., 2019; Wallace, 2016).

Despite the negative association between nestling mitochondrial metabolic rates and the number of nestlings, we did not find any association between nestling ROS production and the number of nestlings. This result suggests that higher metabolism did not lead to higher mitochondrial ROS production in red blood cells in our case. Yet, only 13 individuals raised in small broods were included in ROS production analysis (out of 52), and only 2 juveniles (out of 32), which may explain the lack of association. Furthermore, an increase of mitochondrial metabolism is not always associated with a higher ROS production (see limitations below).

In contrast to our predictions, fledging success was positively associated with the number of nestlings at day 14 (even when excluding the very small broods from the analysis), while we did not find an association of the brood size a few days before fledging with recapture probability as juveniles. One objective of this study was to assess if differences in nestling mitochondrial metabolic phenotype could predict different juvenile recapture probabilities. In our case, we did not find any association of nestling mitochondrial metabolic rates on juvenile apparent survival. We may have expected higher mitochondrial metabolism to lead to detrimental consequences through an increase in ROS release (potentially leading to oxidative stress). However, as previously stated, ROS production did not differ between nestlings and both results are concordant. Furthermore, if nestlings that survived until day 14 were subject to selective disappearance, testing for the association between mitochondrial phenotype and survival as juvenile seems challenging.

As a limitation in our study, mitochondrial ROS production, substrate preferences and mitochondrial aerobic metabolism are known to vary between tissues (Mailloux, 2020; Salmón et al., 2022). Therefore, one should always be careful when investigating ROS production in a single tissue (Costantini, 2019; Monaghan et al., 2009). However, we focused our study on blood samples to i) estimate nestling survival and potential long-lasting effect of our experiment and ii) since mitochondrial aerobic metabolism measurements in blood samples can be positively associated with other tissues (Koch et al., 2021; Stier et al., 2017). Collecting blood samples allows the use of limited-invasive methods on wild species, and to avoid terminal sampling.

Altogether, our results suggest that nestling mitochondrial aerobic metabolism is associated with the actual number of nestlings in the nest, and the contribution of postnatal environmental conditions experienced by the offspring explains a large part of the variation. The effect of rearing conditions on offspring mitochondrial metabolism emphasizes the plasticity of mitochondrial metabolism in changing environments. Further studies would be needed to closely investigate what are the major environmental cues affecting the offspring mitochondrial metabolism during the growth period (e.g., availability of nutrients, ambient temperature) (White & Kearney, 2013), but also to disentangle the role of the brood size in influencing rearing environment (e.g., nest temperature (Andreasson et al., 2016)) and its consequences on nestling physiology and fitness-related traits (e.g., body temperature, DNA methylation, ageing) (Andreasson et al., 2018; Koch et al., 2021; Sheldon et al., 2018).

## Supporting information

Electronic supplementary materials

## Acknowledgements

We are grateful to Toni Laaksonen, Jorma Nurmi, Robin Cristofari, Natacha Garcin, Ida Penttinen, Bin-Yan Hsu and volunteer bird ringers for their help on the field. We thank Tuija Koivisto for the video analysis. We thank Marine Pery for her involvement in this project.

## Ethics

All procedures were approved by the Animal Experiment Committee of the State Provincial Office of Southern Finland (license no. ESAVI/5454/2020) and by the Environmental Center of Southwestern Finland (license no. VARELY/890/2020) granted to S.R.

## Authors contribution

S.R, A.S had the original idea and designed the study with N.C-S. N.C-S, S.R, A.S, M.H collected the data. N.C-S and A.S collected mitochondrial respiration rates measurements. N.C-S performed DNA extractions and conducted qPCR analysis in collaboration with S.Z. N.C-S conducted statistical analysis and wrote the first version of this manuscript under the supervision of S.R and K.A. All co-authors revised the manuscript. S.R, A.S, V.A-V funded experimental work and data collection.

## Data available statement

Data are available on Figshare DOI: 10.6084/m9.figshare.22354432 (embargo pending upon publication).

