## Supplementary material for "Early-life environmental effects on mitochondrial aerobic metabolism: an experimental brood size manipulation in wild great tits": Electronic supplementary materials

Nina Cossin-Sevrin

Department of Biology, 20014 University of Turku, Finland

**A. Complementary analysis on the complete dataset: mitochondrial control flux ratios (FCRs)**

**ESM.Table 1. Results of linear mixed model testing the associations between the number of nestlings in the nest (14 days after hatching) and mitochondrial flux ratios measured on 14-day-old nestlings (N = 102 individuals, n = 55 nest boxes).** Linear mixed models (LMM) estimates are reported with their 95% CI. The original nest box ID and the nest box of rearing ID were both included as random intercepts in the models, except for $FCR_{ROUTINE/CI+II}$ where the original nest box ID could not be included as random intercept because of convergence issue. σ2, within group variance; τ00 between-group variance. Bold indicates significance ($P$ < 0.05).

| | *OXPHOS coupling efficiency* | | | $FCR_{ROUTINE/CI+II}$ | | | $FCR_{CI/CI+II}$ | | |
|---|---|---|---|---|---|---|---|---|---|
| Predictors | Estimates | CI 95% | P-value | Estimates | CI 95% | P-value | Estimates | CI 95% | P-value |
| (Intercept) | 0.93 | 0.89 – 0.97 | **<0.001** | 0.11 | 0.03 - 0.19 | **0.006** | 0.71 | 0.63 – 0.79 | **<0.001** |
| number of nestlings | 8e-4 | -6e-4 – -2e-3 | 0.248 | 1e-3 | -2e-3 – -4e-3 | 0.526 | 1e-3 | -1e-3 – 3e-3 | 0.442 |
| hatching date | -1e-3 | -2e-3 – -3e-4 | **0.004** | 2e-3 | 8e-4 – 3e-3 | **0.002** | -8e-4 | -2e-3 – 6e-4 | 0.266 |
| Random effects | | | | | | | | | |
| σ2 | <0.001 | | | <0.001 | | | <0.001 | | |
| τ00 nest of origin | <0.001 | | | | | | <0.001 | | |
| τ00 nest of rearing | <0.001 | | | <0.001 | | | <0.001 | | |
| Observations | 102 | | | 102 | | | 102 | | |
| Marginal R2 / Conditional R2 | 0.133 / 0.589 | | | 0.152 / 0.489 | | | 0.023 / 0.556 | | |

**B. Results for the control group only**

**ESM.Table 2. Results of linear mixed model testing the associations between the number of nestlings in the nest (14 days after hatching) and mitochondrial respiration rates measured on 14-day-old nestlings (N = 26 individuals from the control group only).** We found similar results as in statistical analysis conducted on the whole data set (same direction for significant effects). Mitochondrial respiration rates were corrected for the mitochondrial DNA copy number (i.e., proxy of mitochondrial density). Linear mixed models (LMM) estimates are reported with their 95% CI. The original nest box ID and the nest box of rearing ID were both included as random intercepts in the models. σ2, within group variance; τ00 between-group variance. Bold indicates significance (*P* < 0.05).

| | *ROUTINE* | | | *CI* | | | *CI + II* | | | *LEAK* | | |
|---|---|---|---|---|---|---|---|---|---|---|---|---|
| Predictors | Estimates | CI 95% | P-value | Estimates | CI 95% | P-value | Estimates | CI 95% | P-value | Estimates | CI 95% | P-value |
| (Intercept) | 6.85 | 1.13 – 12.57 | **0.026** | 27.36 | 12.74 – 41.98 | **0.002** | 42.52 | 18.77 – 66.28 | **0.002** | 4.84 | 2.21 – 7.48 | **0.002** |
| number of nestlings | -0.26 | -0.65 – 0.12 | 0.162 | -1.18 | -2.14 – -0.22 | **0.020** | -1.82 | -3.38 – -0.25 | **0.026** | -0.25 | -0.42 – -0.09 | **0.007** |
| mtDNAcn | 0.29 | -0.03 – 0.60 | 0.070 | 0.54 | -0.26 – 1.34 | 0.175 | 0.88 | -0.39 – 2.16 | 0.163 | 0.14 | 0.01 – 0.26 | **0.037** |
| hatching date | - 0.04 | -0.15 – 0.07 | 0.433 | -0.18 | -0.46 – 0.10 | 0.179 | -0.30 | -0.75 – 0.16 | 0.181 | - 0.03 | -0.08 – 0.02 | 0.252 |
| Random effects | | | | | | | | | | | | |
| σ2 | 0.53 | | | 2.95 | | | 6.58 | | | 0.05 | | |
| τ00 nest of origin | 0.01 | | | 1.47 | | | 3.28 | | | 0.08 | | |
| τ00 nest of rearing | 0.45 | | | 2.56 | | | 8.35 | | | 0.09 | | |
| Observations | 26 | | | 26 | | | 26 | | | 26 | | |
| Marginal R2 / Conditional R2 | 0.483 / 0.724 | | | 0.585 / 0.824 | | | 0.571 / 0.845 | | | 0.684 / 0.932 | | |

**C. Complementary analysis without including the small brood sizes**

**ESM.Table 3. Results of linear mixed model testing the associations between the number of nestlings in the nest (14 days after hatching) and mitochondrial respiration rates measured on 14-day-old nestlings (N = 90 individuals from 46 nests, broods having less than 5 nestlings at day 14 are not included in the analysis).** Mitochondrial respiration rates were corrected for the mitochondrial DNA copy number (i.e. proxy of mitochondrial density). Linear mixed models (LMM) estimates are reported with their 95% CI. The nest box of rearing ID was included as random intercept in the models. Nest box of origin could not be included as random intercept because of convergence issues. σ2, within group variance; τ00 between-group variance. Bold indicates significance (*P* < 0.05).

| | *ROUTINE* | | | *CI* | | | *CI + II* | | | *LEAK* | | |
|---|---|---|---|---|---|---|---|---|---|---|---|---|
| Predictors | Estimates | CI 95% | P-value | Estimates | CI 95% | P-value | Estimates | CI 95% | P-value | Estimates | CI 95% | P-value |
| (Intercept) | 3.84 | 1.53 – 6.16 | **0.002** | 16.69 | 9.32 – 24.07 | **<0.001** | 24.27 | 12.82 – 35.72 | **<0.001** | 2.37 | 1.06 – 3.69 | **0.001** |
| number of nestlings | -0.09 | -0.20 – 0.03 | 0.143 | -0.29 | -0.66 – 0.08 | 0.117 | -0.39 | -0.96 – 0.19 | 0.182 | -0.04 | -0.10 – 0.03 | 0.247 |
| mtDNAcn | 0.30 | 0.20 – 0.40 | **<0.001** | 0.74 | 0.48 – 1.00 | **<0.001** | 1.18 | 0.78 – 1.59 | **<0.001** | 0.15 | 0.10 – 0.20 | **<0.001** |
| hatching date | -0.01 | -0.05 – 0.03 | 0.563 | -0.12 | -0.24 – -0.002 | 0.055 | -0.18 | -0.36 – 0.01 | 0.062 | -0.01 | -0.03 – 0.01 | 0.246 |
| Random effects | | | | | | | | | | | | |
| σ2 | 0.33 | | | 2.09 | | | 4.91 | | | 0.08 | | |
| τ00 nest of rearing | 0.32 | | | 3.91 | | | 9.51 | | | 0.12 | | |
| Observations | 90 | | | 90 | | | 90 | | | 90 | | |
| Marginal R2 / Conditional R2 | 0.293 / 0.643 | | | 0.250 / 0.739 | | | 0.248 / 0.744 | | | 0.248 / 0.682 | | |

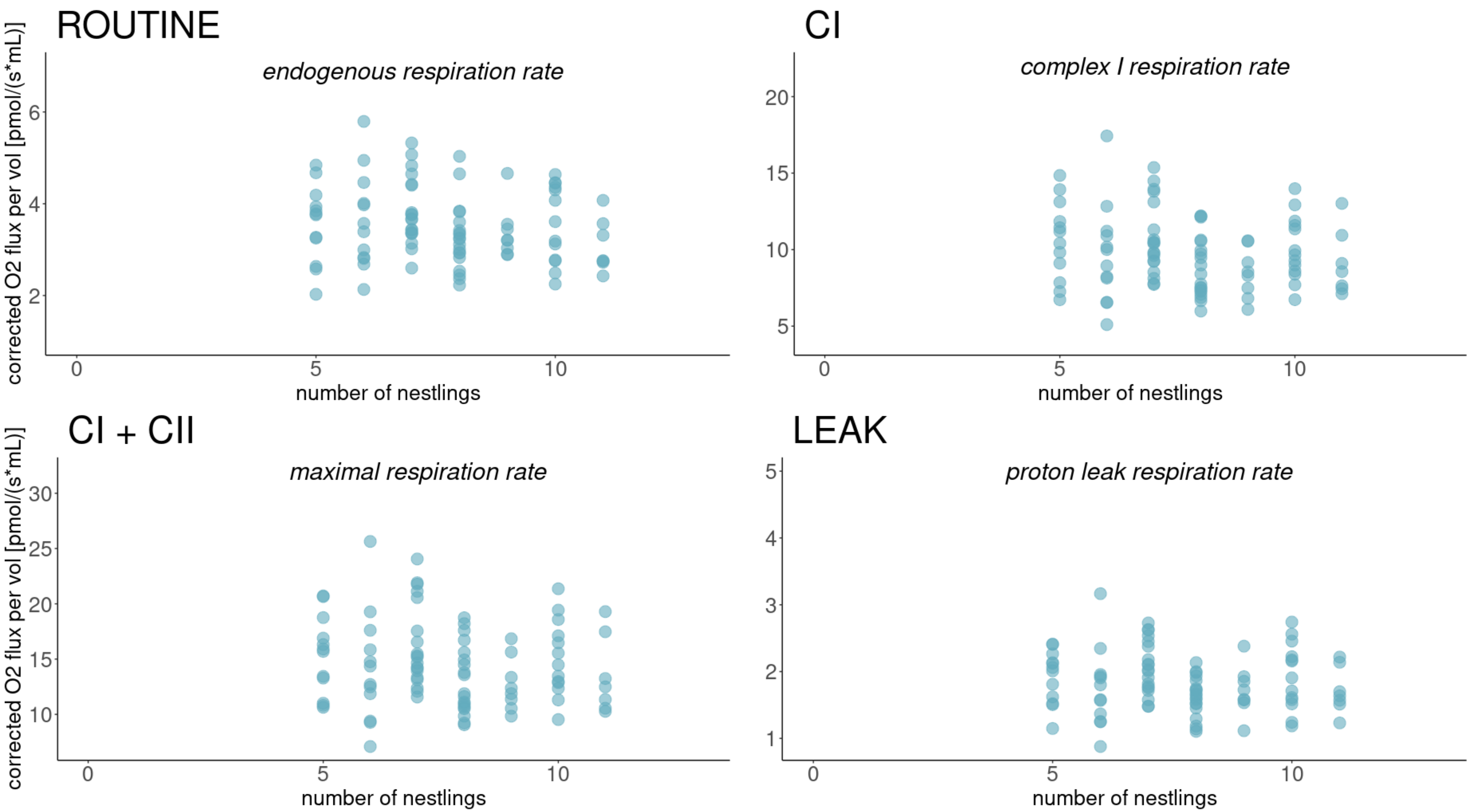


**ESM.Fig.1. Predicted values of mitochondrial respiration rates on 14 days old nestlings according to the number of nestlings in the nest (at day 14).** N = 90 individuals, small broods (< 5 nestlings) are not included in the graph. Predicted values are extracted from linear mixed models (LMMs). Mitochondrial respiration rates were corrected for mitochondrial DNA copy number (i.e., proxy of the mitochondrial density). The original nest box ID and the nest box of rearing ID were both included as random intercepts in the models. R2 of each model are reported in ESM.Table 3.

**D. Results for the ROS production measurements**

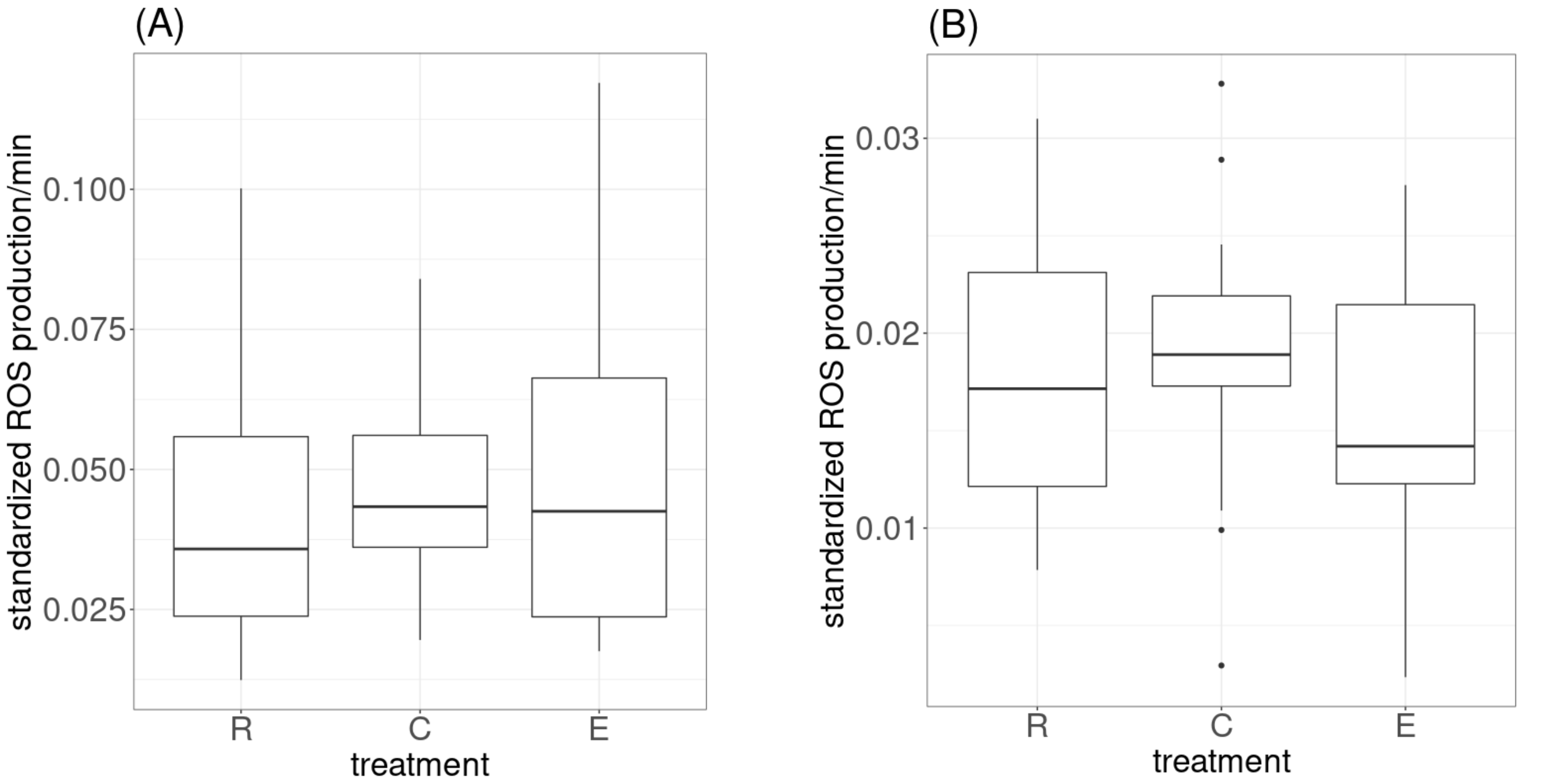


**ESM.Fig.2. Standardized Reactive Oxygen Species (ROS) production per minute according to the treatment groups (R: reduced brood, C: control brood, E: enlarged brood) measured in 14-days-old nestlings (A) and in juveniles (B).** Raw data box plots are presented with their 95% CI. Standardized ROS production per minute was corrected for the number of cells in samples.

**ESM.Table 4. Results of linear mixed model testing the impact of the brood size manipulation treatment groups (R: reduced, C: Control, E: enlarged) on nestling (14 days-old) mitochondrial ROS production (A) and results from linear model testing the impact of the brood size manipulation on juvenile mitochondrial ROS production (B).** Mitochondrial ROS productions were corrected for the mitochondrial DNA copy number (i.e., proxy of mitochondrial density). Linear mixed model and linear model estimates are reported with their 95% CI. The nest box of rearing ID was included as random intercept in the model A, but could not be included as random intercept(s) because of convergence issues. σ2, within group variance; τ00 between-group variance. Bold indicates significance ($P$ < 0.05). See sample-sizes in Table 1.

| | 1. nestling mitochondrial *ROS* production | | | 2. juvenile mitochondrial *ROS* production | | |
|---|---|---|---|---|---|---|
| Predictors | Estimates | CI 95% | P-value | Estimates | CI 95% | P-value |
| (Intercept) | 0.07 | 2e-3 – 0.13 | **0.043** | 0.01 | -3e-4 – 0.02 | 0.057 |
| treatment (E) | 4e-4 | -0.01 – 0.01 | 0.956 | -5e-3 | -9e-3 – -7e-4 | **0.022** |
| treatment (R) | -6e-3 | -0.02 – 0.01 | 0.396 | -4e-3 | -9e-3 – 2e-3 | 0.162 |
| initial brood size | -4e-4 | -4e-3 – 3e-3 | 0.823 | 7e-4 | -7e-4 – 2e-3 | 0.318 |
| mtDNA*cn* | 7e-4 | -1e-3 – 3e-3 | 0.492 | 3e-3 | 3e-3 – 6e-3 | **0.037** |
| hatching date | -3e-4 | -1e-3 – 6e10-4 | 0.490 | | | |
| Random effects | | | | | | |
| σ2 | <0.001 | | | | | |
| τ00 nest of rearing | <0.001 | | | | | |
| Observations | 94 | | | 53 | | |
| Marginal R2 / Conditional R2 | 0.033 / 0.617 | | | 0.151/ 0.080 | | |

**E. Results for the association between mitochondrial respiration rates in 14-days-old nestlings and survival as juvenile**

**ESM. Table 5. Results of a GLMM (with logistic binary distributions of the dependent variables, survival: 0=dead, 1=alive) testing whether mitochondrial respiration rates measured at day 14 predict juvenile recapture probability (i.e., proxy of medium-term apparent survival).** Models only include individuals for which mitochondrial metabolic rates have been measured at day 14 (N = 102 individuals). 67 individuals (from 34 nests) have been recaptured as juveniles. Random intercepts could not be included in the models because of convergence issues. Odds ratios are reported with their 95% CI. R2 values are estimated from the *coefficient of determination D* (Tjur's approach). Bold indicates significance (*P* < 0.05).

| | *ROUTINE* | | | *CI* | | | *CI+II* | | | *LEAK* | | |
|---|---|---|---|---|---|---|---|---|---|---|---|---|
| Predictors | Odds Ratios | CI 95% | P-value | Odds Ratios | CI 95% | P-value | Odds Ratios | CI 95% | P-value | Odds Ratios | CI 95% | P-value |
| (Intercept) | 82.24 | 0.43 – 1.7e4 | 0.10 | 60.96 | 0.26 – 1.4e4 | 0.14 | 75.12 | 0.35 – 1.7e4 | 0.11 | 114.31 | 0.67 – 2.2e4 | 0.07 |
| Mitochondrial respiration rate d14 | 1.30 | 0.87 – 1.95 | 0.20 | 1.08 | 0.95 – 1.23 | 0.24 | 1.04 | 0.96 – 1.14 | 0.31 | 1.33 | 0.66 – 2.47 | 0.38 |
| hatching date | 0.89 | 0.80 – 0.97 | **0.01** | 0.89 | 0.81 – 0.98 | **0.01** | 0.89 | 0.80 – 0.97 | **0.01** | 0.89 | 0.80 – 0.98 | **0.01** |
| Observations | 102 | | | 102 | | | 102 | | | 102 | | |
| R2 | 0.09 | | | 0.09 | | | 0.08 | | | 0.08 | | |
